# Improved growth and morphological plasticity of *Haloferax volcanii*

**DOI:** 10.1101/2020.05.04.078048

**Authors:** Roshali T. de Silva, Mohd F. Abdul-Halim, Dorothea A. Pittrich, Hannah J. Brown, Mechthild Pohlschroder, Iain G. Duggin

## Abstract

Some microbes display pleomorphism, showing variable cell shapes in a single culture, whereas others differentiate to adapt to changed environmental conditions. The pleomorphic archaeon *Haloferax volcanii* commonly forms discoid-shaped (‘plate’) cells in culture, but may also be present as rods, and can develop into motile rods in soft agar, or longer filaments in certain biofilms. Here we report improvement of *H. volcanii* growth in both semi-defined and complex media by supplementing with eight trace-element micronutrients. With these supplemented media, transient development of plate cells into uniformly-shaped rods was clearly observed during the early log phase of growth; cells then reverted to plates for the late log and stationary phases. In media prepared with high-purity water and reagents, without supplemental trace elements, rods and other complex elongated morphologies (‘pleomorphic rods’) were observed at all growth stages of the culture; the highly-elongated cells sometimes displayed a substantial tubule at one or less frequently both poles, as well as unusual tapered and highly-curved forms. Polar tubules were observed forming by initial mid-cell narrowing or tubulation, causing a dumbbell-like shape, followed by cell division towards one end. Formation of the uniform early-log rods, as well as the pleomorphic rods and tubules were dependent on the function of the tubulin-like cytoskeletal protein, CetZ1. Our results have revealed the remarkable morphological plasticity of *H. volcanii* cells in response to multiple culture conditions, and should facilitate the use of this species in further studies of archaeal biology.

## Introduction

Archaea exhibit some of the most diverse and unusual microbial cell morphologies. Cell shapes range from rods and cocci to striking triangles and squares. However, little is known about the mechanisms and environmental cues that dictate archaeal cell morphology, or the regulation of transitions between cell morphologies. The model haloarchaeon, *Haloferax volcanii*, when first isolated, was described as mainly disk-shaped cells, with cell shape and size varying significantly (1). In routine liquid cultures of *H. volcanii*, both pleomorphic discoid (plate) and rod cell morphologies may be observed, occasionally with more unusual shapes (2–4).

The conditions that influence *H. volcanii* cell shapes and the relative abundance of the distinct types are not well understood, and specific signals have not been identified. In *H. volcanii* biofilms, substantial elongation (filamentation) has been observed in a subpopulation of cells (5). Furthermore, *H. volcanii* forms motile rods in soft-agar, where rods are observed at the forefront of expanding colonies of swimming cells (6). The tubulin-like cytoskeletal protein CetZ1 is required for rod-formation, suggesting a connection between morphology and motility that may be expected based on improved hydrodynamics or directional movement of rods (4, 6). Conversely, the peptidase archaeosortase A (ArtA) and phosphatidylethanolamine biosynthesis enzymes PssA and PssD, which are required for the C-terminal processing and covalent lipid attachment of several *H. volcanii* surface proteins including the surface (S-layer) glycoprotein, are required for effective and stable plate-shaped cell formation (7, 8).

To better understand cell shape switching and the significance of the distinct shapes, we aimed to define conditions that can be robustly used to study these processes. We determined that inclusion of a trace elements (TE) solution in both complex and semi-defined culture media substantially improves culture and cell shape reproducibility for this species. In the new media, early-log rod formation was also more reproducible, while the pleomorphic rods and other shapes seen in cultures lacking TE were absent. We further defined culture conditions and characterized these CetZ1-dependent morphological changes during both TE-depletion and early-log rod formation in replete cultures. These conditions may serve as experimental models for studies of morphological development in archaea.

## Methods

### Archaeal strains

*Haloferax volcanii* H26 (Δ*pyrE2*), H53 (Δ*pyrE2* Δ*trpA*) or H98 (Δ*pyrE2* Δ*hdrB*) (7), containing the pHV2-based plasmids pTA962 or pTA963 (9) as indicated, were used as wild-type strains. *H. volcanii* ID59 (6), transformed with pTA962 (i.e. ID113), was the *cetZ1* knockout strain used (based on H98). Attempts to verify the roles of CetZ1 by plasmid-based expression of wild-type *cetZ1* in the Δ*cetZ1* strain (ID59) were unsuccessful, due to polar effects on a downstream gene (unpublished results), so we expressed *cetZ1.E218A* (a dominant-inhibitory mutant) to confirm the involvement of *cetZ1*, by using H98 + pTA962-*cetZ1.E218A*, as described previously (6).

### Culture media and general growth conditions

Media were based on the Hv-Ca and Hv-YPC media commonly used for *H. volcanii* (10). In this study, we used defined sources of all media reagents, which we found to be important for reproducibility. However, the data reporting the initial observation of early-culture rod development (Fig. 1) were obtained with the original recipe for Hv-Ca (10). For solid media, Bacteriological Agar (Oxoid LP0011) (10 g/L), was dissolved in the volume of water required for the medium by heating in a microwave oven, before mixing with pre-heated concentrated stock solutions of macronutrients and buffered salt water solutions (listed below) and autoclaving, followed by addition of trace elements or heat-sensitive reagents. Water was obtained from an ultrapure (18.2 MOhm.cm) water purification system (Sartorius).

**Figure 1.**
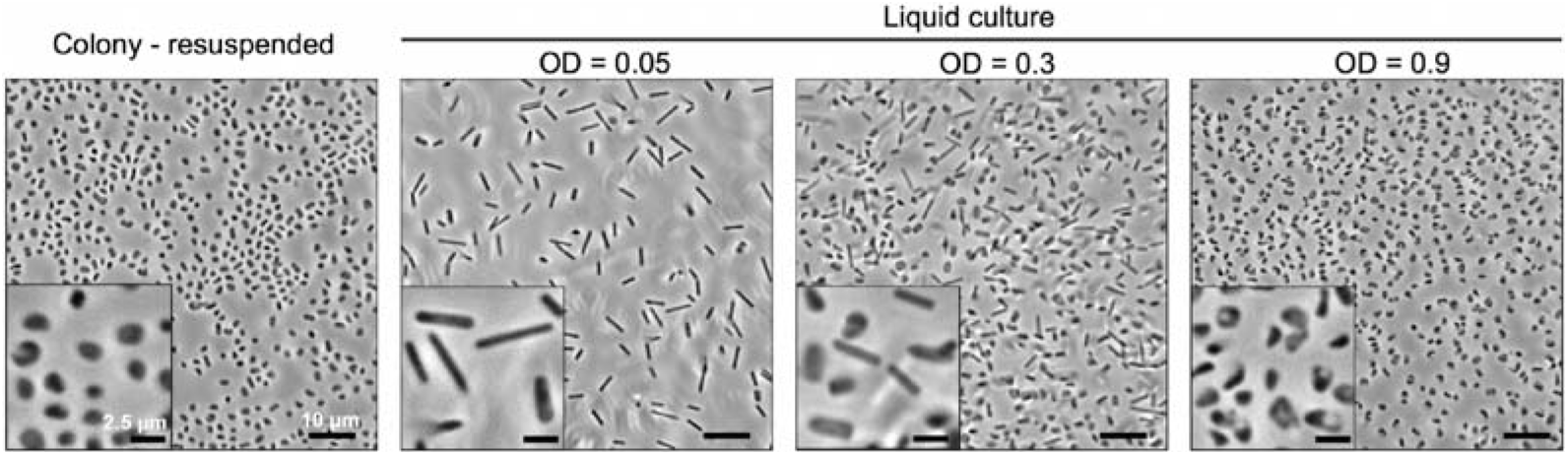
*H. volcanii* forms rods during the early stages of growth at low culture density in liquid batch culture. A fresh colony of *H. volcanii* (H53 + pTA963) was used to inoculate Hv-Ca liquid medium (without additional vitamins or trace elements, but prepared with standard-purity reagents as the only possible source of trace elements). Cells from early-, mid- and late-log phase cultures (OD_600_ 0.05, 0.3, and 0.9, respectively) were observed by phase-contrast microscopy. Scale bars are 10 μm (inset = 2.5 μm).

Hv-Ca medium was prepared similarly to that described previously (10), however, in all experiments except Fig. 1, Hv-Ca medium also contained an additional vitamin solution. Hv-Ca contained: 144 g/L NaCl (Sigma S6191), 21 g/L MgSO_4_•7H_2_O (Sigma M1880), 18 g/L MgCl_2_•6H_2_O (Sigma M2393), 4.2 g/L KCl (Sigma P5405), 12 mM Tris-HCl (pH 7.4) (Sigma RDD008, buffered with AR grade HCl), 3 mM CaCl_2_ (Sigma C5670), and 5 g/L Casamino Acids (Oxoid LP0041, prepared initially as a 10x stock solution). The medium was then autoclaved. Once cool, a filter-sterilized vitamin concentrated stock solution (1000x), containing 1 g/L thiamine (Sigma T1270) and 0.1 g/L biotin (Sigma B4639), was added (1 mL per L of medium). Buffered salt water (BSW) was prepared as the buffered salts solution described above, i.e. Hv-Ca medium (18 % w/v total salts), omitting the Casamino acids and vitamins. BSW was prepared as a concentrated (30% w/v salts) stock solution.

To prepare Hv-Cab medium, a trace-elements concentrated stock solution (100x) was added (10 mL per L) to the Hv-Ca medium after autoclaving and cooling (including vitamins). The 100x trace-elements stock solution contained: 5 g/L Na_2_EDTA•2H_2_O (Sigma E1644), 0.8 g/L FeCl_3_ (Sigma 157740), 0.05 g/L ZnCl_2_ (Sigma 793523), 0.01 g/L CuCl_2_ (Sigma 751944), 0.01 g/L CoCl_2_ (Sigma 232696), 0.01 g/L H_3_BO_3_ (Sigma B6768), 1.6 g/L MnCl_2_ (Sigma 328146), 0.01 g/L NiSO_4_ (Sigma 656895), and 0.01 g/L Na_2_MoO_4_•2H_2_O (Sigma M1003); the pH of the solution was adjusted to 7.0 with NaOH (AR grade, Sigma), and then the obvious brown precipitate that formed on neutralization was removed by 0.2 μm sterile filtration; stock solutions were stored at room temperature in sterile aliquots.

Hv-YPC medium (10) was prepared as per the Hv-Ca medium above, except the supplementary vitamins were omitted, the Casamino acids was used at a concentration of 1 g/L, and 5 g/L Yeast Extract (Oxoid LP0021) and 1 g/L Peptone (Oxoid LP0037) were included (initially dissolved as a 10x stock solution). To prepare “Hv-YPCab” medium, the 100x trace-elements stock solution was added (10 mL per L of medium) to Hv-YPC after autoclaving and cooling.

Most cultures (5 mL) were grown in sterile 50 mL Falcon tubes, incubated at 45°C with shaking at 200 rpm in a GFL 1092 rotary-shaking water bath (the tube’s lid was maintained loosened) unless otherwise indicated. *H. volcanii* was initially grown on solid medium followed by continuous growth in liquid medium (diluted once per day), to maintain steady logarithmic growth prior to sampling, which was only done after the culture had been growing steadily in continuous log-phase for at least 2 days, unless otherwise indicated. For obtaining growth curves in microtitre plates, cultures established as above at 45°C were used to prepare a culture with a starting OD_600_ of 0.1 in the indicated Hv-YPC based media, with or without TEs as indicated. Growth at 42°C with 200 rpm shaking was then monitored in a Tecan Spark microtiter plate spectrophotometer (30 min reading time intervals).

### Time-course studies of rod development

In order to obtain a standardized way of observing rod cells, and analyzing rod-development, time-course studies of the early stages of liquid culture growth were carried out (Fig. 4). The *H. volcanii* strain was streaked onto an agar plate made from the same medium as chosen for the assay (*e.g.* Hv-YPCab), and then the plate was incubated in sealed humidified bag at 45**°**C for 4 days. Colonies were collected from the surface with a microbiological loop, and resuspended in 5 mL of liquid medium, sufficient to give an OD_600_ > 0.05. The OD_600_ was immediately adjusted to 0.05 by dilution with fresh pre-warmed medium, marking the start of the time-course. Samples were withdrawn as required for microscopy or other analyses (Fig. 4). In some experiments, cells were obtained directly from a mid-log liquid culture (Fig. S1B).

To observe trace element limitation over a time-course (Fig. 4), cultures were first grown into steady log phase in Hv-Cab medium at 45°C, with shaking at 200 rpm (5 mL culture). A sample of the culture was withdrawn and centrifuged at 5,000 rpm for 5 min in an Eppendorf microcentrifuge. The supernatant was discarded and the cells were then washed by re-suspending the pellet in 1 mL of 18% BSW. The wash was repeated three times and then the cells were finally re-suspended in 0.1 mL 18% BSW and then spread onto Hv-Ca agar plate(s). The plate(s) were placed into a plastic bag and incubated at 45°C for 4 days. 5 mL of liquid Hv-Ca was added to the plate surface and the cells were re-suspended, avoiding disturbing the agar. The OD_600_ was adjusted to 0.05 with Hv-Ca medium, to start the time course (incubation was continued at 45°C at 200 rpm).

### Microscopy

Samples were prepared by mounting a 2 μl volume of culture (concentrated where necessary by centrifugation and gentle resuspension) onto a slide prepared with a ~170 μm thick, 1% agarose pad containing 18% BSW (6). Most phase-contrast images were acquired with a Zeiss AxioPlan2, Nikon Ti, or a V3 Delta Vision Elite (Applied Precision Inc., a GE Healthcare Company, Issaquah, USA) microscope, with 1.4 NA phase-contrast objectives. Images were analyzed using the MicrobeJ plug-in for ImageJ (11). The cell circularity (4π × area / perimeter^2^) was measured on the interpolated contour generated using the particle medial axis and the specified value of the width measured along the medial axis. Circularity values range between 0 and 1. Cell elongation was calculated as the inverse of the circularity.

For live cell imaging, a CellASIC ONIX microfluidics system (with B04A-03 plates), was used with a Nikon Ti inverted microscope equipped with an incubated stage (42°C). The four flow chambers were initially washed with 1 mg/mL Bovine Serum Albumin (BSA) in phosphate-buffered saline followed by rinsing with 18% BSW at 5 psi for 5 min. Cells of *H. volcanii* (H98 + pTA962), freshly cultured on the agar medium indicated with each movie, were resuspended in the respective liquid medium (pre-warmed to 42°C), and then loaded into the chambers as per the manufacturer’s instructions, and the chambers were perfused with medium at 2 psi for 20 h, with time-lapse images recorded at 10 min intervals.

### Coulter cytometry

Culture samples were diluted (1:1000) with 18% BSW, and then analyzed with a Multisizer 4 Coulter cytometer (Beckman-Coulter) as described previously (6).

### Western blotting

Western blotting was carried out as described previously (6).

## Results

It was previously shown that, in soft agar assays of cellular swimming motility, *H. volcanii* cells change from a discoid (plate) shape into a rod shape in a CetZ1-dependent manner (6). However, the soft-agar motility assay is not a robust method for ongoing studies of cell-shape differentiation in *H. volcanii* as the soft agar is difficult to separate from the cells, cell yields are low, and the distribution of shapes observed is sensitive to the site of sampling from the soft agar (6). We and others have noticed that *H. volcanii* can form regularly-shaped rods in the early stages of culture, from around the time the culture transitions from lag to log phase, and then cells revert to the irregular discoidal ‘plate’ cells later in log phase (12). As shown in Fig. 1, cells obtained from colonies grown on agar medium are initially plate-shaped upon resuspension in fresh Hv-Ca liquid medium, but after overnight shaking incubation (to an OD_600_ of 0.05) cells were predominantly rod-shaped. With further incubation, until the culture reached OD_600_ of 0.3, the rod-shaped cells had started to disappear and by 0.9 the culture predominantly contained the plate-shaped cells (Fig. 1). Early culture cells had similar morphology to motile cells withdrawn from motility agar (6), and were recently shown to be motile and suitable for direct observation of swimming (12).

The OD_600_ where rods could be observed and then returned to plates varied between cultures, possibly due to differences in the size of starting inoculum and the times cultures take to reach equal culture density. We have also noticed widely varying growth yields of *H. volcanii* in Hv-Ca medium prepared in different laboratories or with different sources of reagents, and the extent and timing of early-culture rod formation is generally sensitive to different media sources (e.g. we observed that one lower-grade source of NaCl inhibited the early-culture rod development that was seen in all other media tested). These issues make reproducing certain experiments difficult between laboratories using different reagents. Such considerations are pertinent when nutrition, metabolism, stress-response or whole-organism (‘omic’) projects are planned.

The supply of trace elements in the medium was also observed to have dramatic effects on growth and morphology, as described further below. The results shown in Fig. 1 were obtained with Hv-Ca medium without supplemental trace elements, but growth in this culture was satisfactory as cells must have had sufficient supply of essential trace elements in this medium. However, such growth was not always observed with other reagent sources. We therefore sought to understand and standardize the conditions required for optimal growth and rod development; this led to the identification of previously uncharacterized morphological responses of *H. volcanii.*

### H. volcanii growth and morphology in nutrient-limited batch culture

The commonly used media for *H. volcanii*, Hv-Ca and Hv-YPC, consist of macronutrients for heterotrophic growth in a base of 18% (w/v) buffered primary sea-water salts (10). Hv-Ca is a semi-defined medium with Casamino acids as the carbon and energy source, and typically provides a slower growth rate and yield than the complex Hv-YPC medium, which contains Yeast extract, Peptone and Casamino acids added to the 18% salts base. Both media are known to produce different morphotypes in the same culture—primarily a mix of rods and plate cells, with some other more unusual shapes also sometimes seen.

Since these media are normally not supplemented with micronutrients, beyond what may be present as contaminants in the media reagents (10), and a range of metal ions or other elements are typically required by cells at low or moderate concentrations to act as essential cofactors in enzymatic reactions, we added a solution of eight compounds as a source of micronutrients to the Hv-Ca medium. This was based on common requirements for microorganisms, and contained the elements Fe, Zn, Cu, Co, Mn, Ni, Mo and B (‘trace elements’, TE; see Methods). A vitamin solution (10) was also added. The new medium (named “Hv-Cab”, for the ‘b’ variant of Hv-Ca) substantially improved the growth rate and yield, and their reproducibility, in *H. volcanii* cultures, which was due primarily to the inclusion of the TE solution, as described below.

To identify any growth or morphological changes associated with omission of the primary nutrient sources in Hv-Cab, mid-log *H. volcanii* cells were transferred to specific nutrient-omitted media, which were prepared with high-purity reagents to minimize contaminating sources of trace elements or substances that might inhibit growth or development. Growth and cell morphology were monitored over three successive passages (by dilution into fresh media) under each of these nutrient-omitted conditions. Transfer to media without the carbon and energy source, Casamino acids (CAA), caused cells to rapidly cease growth, as expected (Fig. 2A, maroon data-points). Exclusion of the vitamins had a minor effect on the maximum OD_600_ that was consistent throughout the three successive culture passages, but had no effect on cell shape (Fig. 2A-C). However, cultures without supplementary TE showed successively greater reductions in growth rate and yield with each passage, such that by the third passage growth was very poor, and the culture barely reached a maximal OD_600_ of ~0.3 (Fig. 2C).

**Figure 2.**
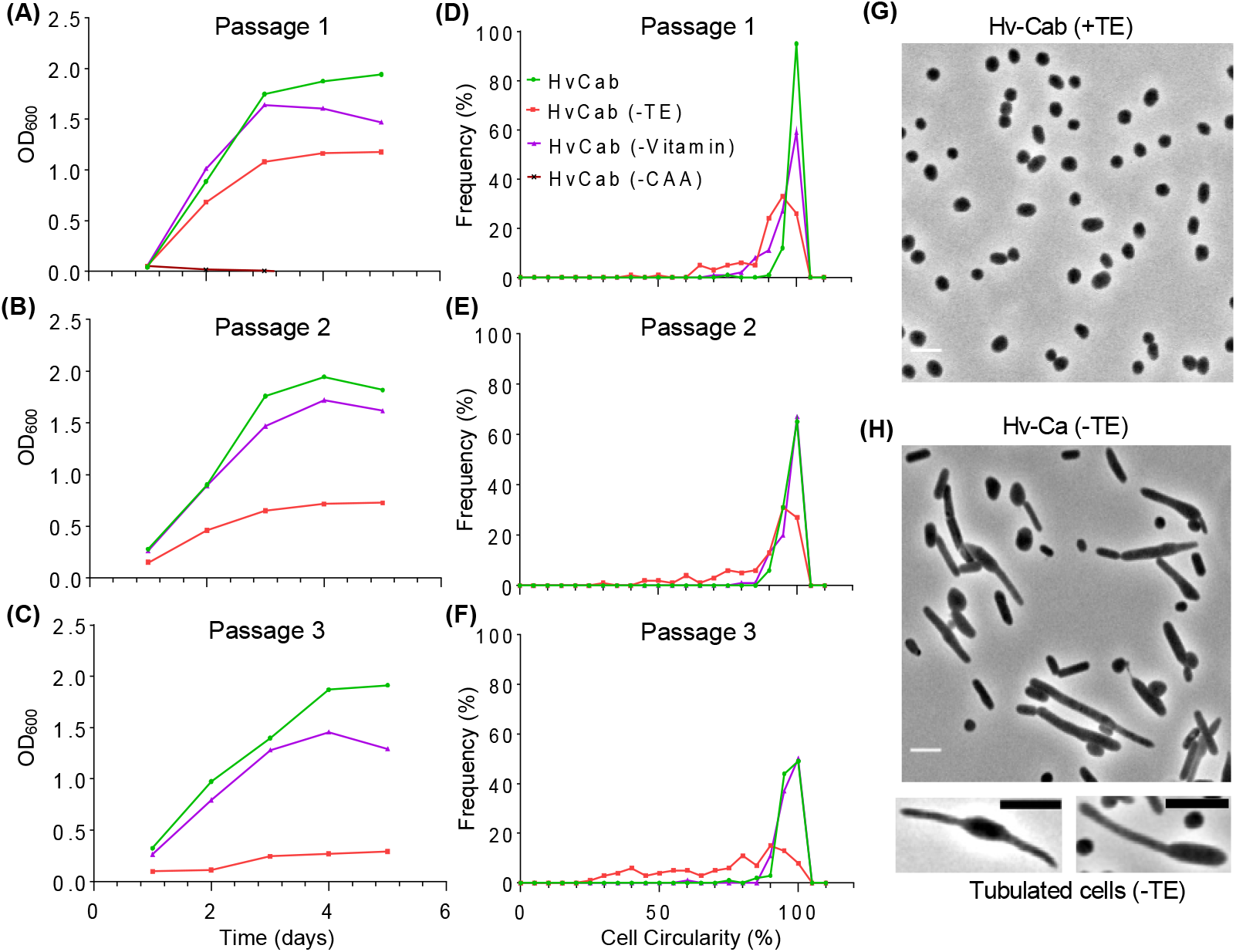
Effects of nutrient depletion on *H. volcanii* growth and morphology in liquid culture. Strain H98 + pTA962 was initially grown to mid-log phase in Hv-Cab (OD_600_ = 0.5), and then was washed 3 times by centrifugation at 5,000 rpm in a microcentrifuge for 5 min, followed by resuspension of the cell pellet in same volume of 18% BSW, and diluted (1:100) into fresh Hv-Cab or in media excluded for one of the major nutrient components, *i.e.* trace elements (TE, red data-points), vitamins (biotin and thiamine, purple triangles), or Casamino acids (CAA, maroon triangles). **(A)** Culture growth was monitored by measuring the OD_600_ over 5-days. Cells from day-5 of the first dilution were then diluted (1:100) into the respective nutrient-depleted media and incubated and monitored the same way for a further 5 days **(B)**. Cells from the second dilution (day-5) were again diluted (1:100) into the respective nutrient-depleted media and then incubated for five more days **(C)**. Cells imaged by phase-contrast microscopy at day-5 (final data point) in each of the cultures (A-C) by determining the circularity of cell outlines (N = 100, randomly selected cells), shown as a corresponding histogram (D-F). G and H shows the phase-contrast images of cells in (G) Hv-Cab (+TE) and (H) Hv-Ca (-TE) on day-5 of the third passage (a representative image of the cells analysed in histogram F). TE depletion also resulted in formation of polar tubules in some cells. The scale bars are 5 μm.

Cells were observed by phase-contrast microscopy at day 5 (stationary phase) after all three of the culture passages, and the cell shapes were visualized and analyzed by automated measurement of the circularity of cell outlines; these data were then plotted as histograms of cell circularity, expressed as a percentage compared to a perfect circle (Fig. 2D-F). The stationary phase Hv-Cab cultures contained almost exclusively plate-shaped cells that had near-circular cell outlines, similar to late log cultures. However, the TE-depleted cultures showed striking cell elongation, which became more extensive in each successive passage (Fig. 2H). Cells depleted of TE showed complex features accompanying the extensively elongated rod shapes, including rods that showed variable widths along their lengths, and some cells exhibiting substantial polar tubules of variable length and width (Fig. 2H).

### Improved growth of H. volcanii in TE-supplemented complex and semi-defined media

Previous studies have shown that in complex medium, such as Hv-YPC (10), elongated and irregular cell morphologies can exist together with the plate-shaped cells (1, 3). Trace elements are expected to be present in Hv-YPC, originating primarily from the Yeast extract. We next investigated the possibility that trace elements may yet be somewhat limiting for *H. volcanii* in Hv-YPC, giving rise to the elongated forms seen in this medium, by supplementing Hv-YPC with the TE solution. By analogy to the Hv-Cab media terminology introduced above, we refer to the TE-supplemented version of Hv-YPC medium as Hv-YPCab.

We initially took mid-log and stationary-phase cells grown in Hv-Cab medium, and then washed and resuspended cells from both starters in Hv-YPC and Hv-YPCab, and then monitored growth in microtiter-plates, to observe how well the cultures grew under these new conditions. The growth rate and yields were noticeably greater in Hv-YPCab medium compared to Hv-YPC (Fig. 3A). The Hv-YPCab culture grew well even when started by inoculation with a stationary phase culture, compared to the Hv-YPC culture inoculated with a mid-log starter, suggesting that the maintenance of supplemental trace elements in the media is useful for minimizing the lag phase and maximizing growth rates and yields of *H. volcanii*.

**Figure 3.**
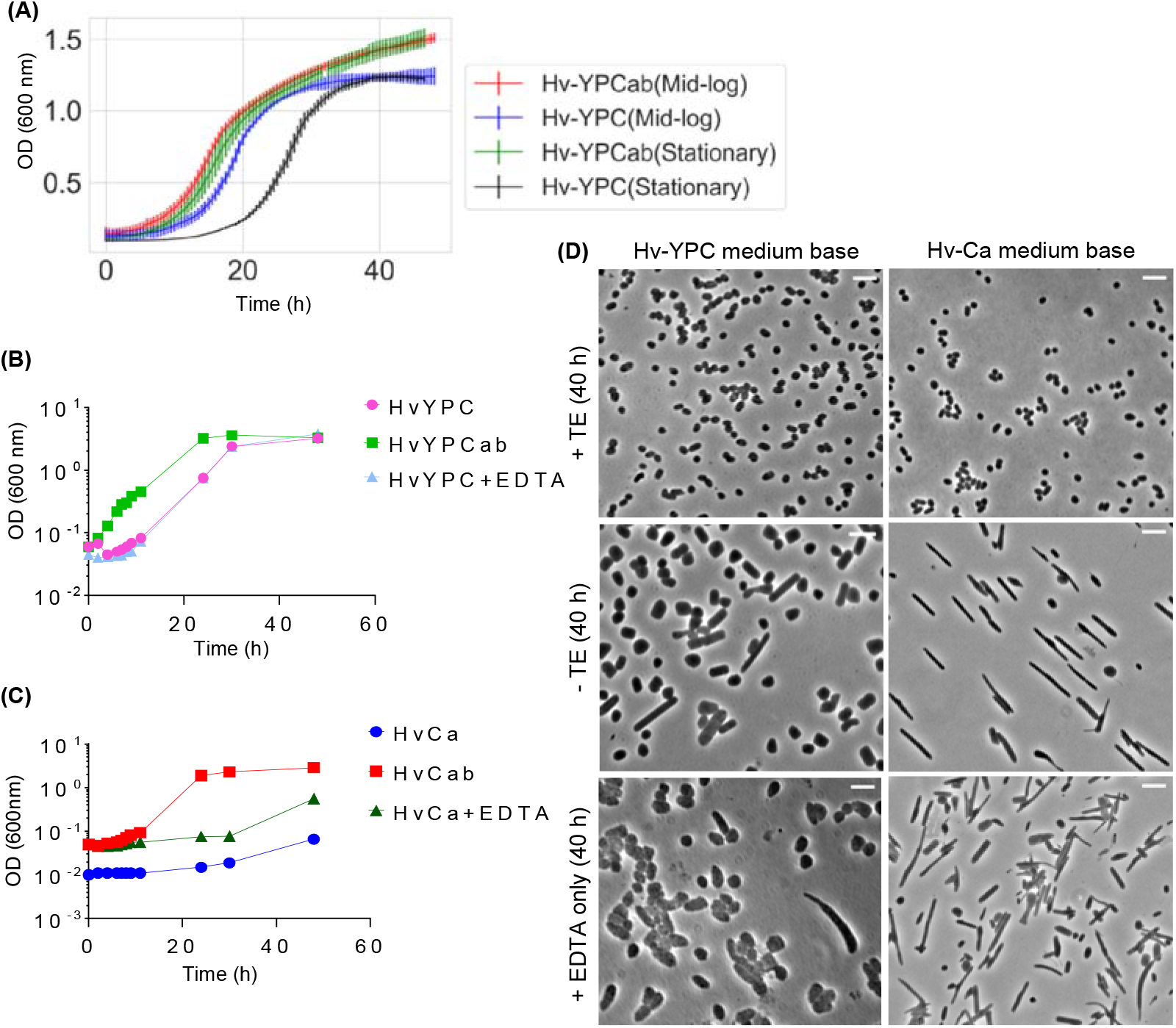
Media supplementation with a trace elements (TE) solution improves growth and cell shape uniformity in *H. volcanii*. (A) Growth curves in microtiter plate format. Two *H. volcanii* (H98 + pTA962) Hv-Cab batch cultures (5 mL) were started from separate colonies. The cells were taken at the mid-log phase (for 6 technical replicates) of one culture and at the stationary phase (for 3 technical replicates) of the other. The error bars indicate the standard deviation. **(B)** Growth curves in tube-culture format, started from steady mid-log cultures in the respective media, comparing Hv-YPC medium with the same media supplemented with the complete TE solution (*i.e.* Hv-YPCab), or with only the EDTA component of the solution (control). **(C)** Tube-culture experiments, as per panel (B) except based on Hv-Ca medium. The Hv-Ca culture was started at ~10-fold lower cell density than the other two, owing to the relatively poor growth of the starter culture. **(D)** Cells sampled after 40 h (stationary phase) were imaged by phase-contrast microscopy. All scale bars represent 5 μm.

**Figure 4.**
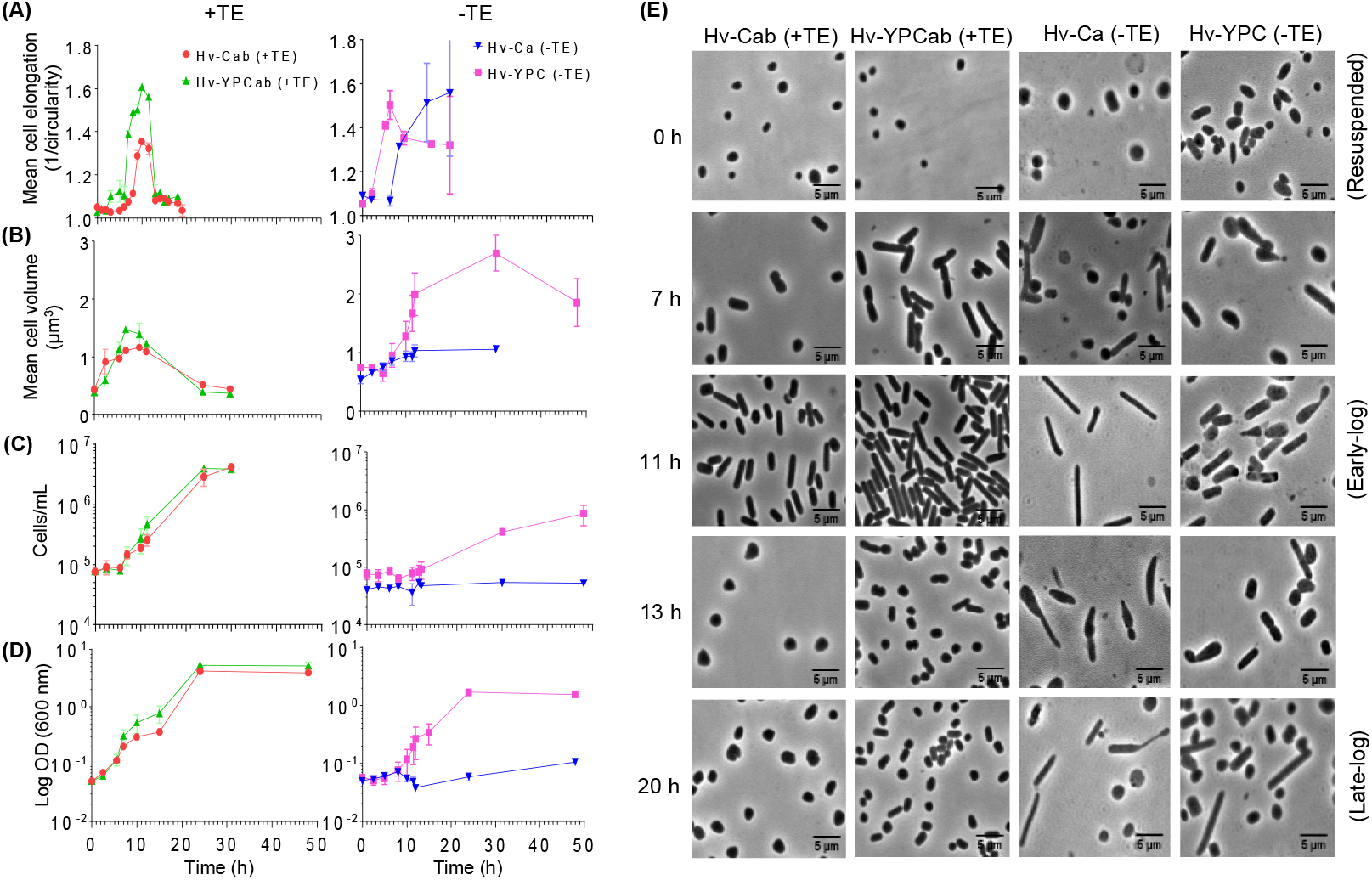
Assays for *H. volcanii* reversible morphological transitions during the growth cycle. Samples of *H. volcanii* (H98 + pTA962), grown in the indicated liquid media (after inoculation from colonies obtained from 4 days of growth on the equivalent solid agar medium), were withdrawn at time-points for microscopy (A)(E), Coulter cytometry determination of cell volumes (B) and counts (C), and OD_600_ measurements (D). Mid-log growth rates (μ μmean±SD) were: Hv-Ca = 0.045±0.011 h^−^ 1, Hv-Cab = 0.149±0.020 h^−1^, Hv-YPC = 0.141±0.008 h^−1^, and Hv-YPCab = 0.206±0.003 h^−1^. The microscopy analysis in panel (A) shows mean cell elongation (expressed as inverse circularity; a value of 1 is a perfect circle and becomes greater as cells become more elongated) of 250 randomly selected cells at each time point. Datapoints shown represent the mean from two independent replicates, and error bars indicate standard deviation of this mean (some are within the datapoints).

We also monitored growth and morphology of cells in standard tube-based batch-cultures. Consistent with the results of Fig. 3A, cultures recommenced growth substantially faster with Hv-YPCab medium, compared to the Hv-YPC medium (Fig. 3B). Similar trends were obtained with experiments based on the Hv-Ca medium (Fig. 3C), although growth was very poor, which was attributed to the lack of trace elements in this medium made with high-purity reagents. Microscopy of late-stage culture samples (at 40 h) revealed that cells were almost uniformly the plate morphotype in Hv-YPCab and Hv-Cab (Fig 3D, ‘+TE’ panels), and showed none of the elongated rods or filaments with tubules previously noted in non-TE-supplemented medium (Fig. 3D, -TE panels). The cell size also appeared substantially smaller in both TE-supplemented media compared to the original media (Fig. 3D). An additional control culture, supplemented with the EDTA only showed that culture growth, cell size and morphology were unaffected compared to Hv-YPC alone, demonstrating that the cells were reacting specifically to the concentration of the trace-element compounds that differ between the original and the supplemented media (Fig 3D). Finally, we also observed a substantial difference in the morphology when comparing cells in Hv-YPC and Hv-Ca (-TE cultures, Fig. 3D); the more elongated cells observed in the Hv-Ca culture were attributed to the more severe TE starvation in this medium (see Fig. 2).

### Development of H. volcanii rods during an early stage of growth in culture

Based on the above results, the new supplemented media were expected to improve consistency and reproducibility of the motile-rod development that occurs during early-culture. It was expected that the elongated/tubulated rod cell types specifically seen during TE-limitation would be absent, thus improving uniformity of the rods seen in early-culture growth. In order to thoroughly characterize the cell morphology development of *H. volcanii* during the batch-culture growth cycle with the new media, we inoculated cultures to a fixed cell density from consistently-prepared starter cultures, and then monitored culture growth rate and cell morphology at regular timepoints. *H. volcanii* was initially cultured for four days on the appropriate solid media to allow colony development, and then sufficient colonies were picked and resuspended in the equivalent liquid medium, and the OD_600_ was adjusted to 0.05 to start the culture at a measurable and reproducible density. Culture growth (OD_600_ and cell number), cell volumes (Coulter cytometry) and morphology (microscopy) were monitored over time, both with or without supplemental TE (Fig. 4).

In both Hv-Cab and Hv-YPCab media, cell volume and culture growth (OD_600_) increased from the start of the experiments (Fig. 4B, 4D) and the cell number began to noticeably increase after ~6 h delay (Fig. 4C). This corresponded approximately to the onset of rod development, as seen in the representative images of the cells (Fig. 4E), and the quantitative analysis of cell elongation (Fig. 4A); in both media (+TE), a peak of rod development was observed commencing ~7-8 h after inoculation, reaching a maximum mean cell elongation after ~10 h, and then from 13 h onwards, plate cells were again almost exclusively seen (Fig. 4a, +TE). Similar results were obtained with cultures of the H26 (+ pTA962) strain inoculated from either agar-based colonies or late-stage liquid cultures (Fig. S1). We also found that early-culture rod development required inclusion of the plasmid, which is based on the natural pHV2 plasmid from *H. volcanii* and complements the uracil-auxotrophy of H26, indicating the importance of plasmid inclusion or prototrophy in these assays (Fig. S1). Comparison of results obtained with the different media showed that the peak of rod development was greater in Hv-YPCab than in Hv-Cab, indicating a higher proportion or longer rods in Hv-YPCab (compare the left-hand two panels in Fig. 4E, 11 h). The timing of rod development during the first few generations and subsequent conversion back to plates approximately corresponded to the timing of an increase (growth) and then a decrease (division) in the mean cell volume, respectively (Fig. 4B).

The effects of TE-starvation during the above colony-resuspension assay were also investigated. Cultures were first grown into mid-log phase in either Hv-Cab or Hv-YPCab medium (*i.e.* +TE), and then they were separated from the medium by centrifugation, washed (media -TE) and plated onto the respective -TE agar medium (Hv-Ca and Hv-YPC). After growth on the agar, cells were resuspended in -TE liquid medium. Cells resuspended from colonies were initially plate-shaped (Fig. 4A, 4E - TE), even though there was a clear limitation of trace elements in the agar, detected by very poor growth of colonies on the Hv-Ca agar. As expected, the Hv-Ca (-TE) liquid cultures showed very poor overall growth, whereas Hv-YPC (-TE) cultures grew quite well (Fig. 4C, 4D). Cells became noticeably elongated by 5 h (Hv-YPC, - TE) or 8 h (Hv-Ca, -TE) of incubation, and they continued to show substantial elongated pleomorphology and increased cell volume for at least 20 h, particularly in the severe starvation conditions (Hv-Ca, -TE) (Fig. 4A, 4B, 4E).

Thus, as may be expected based on the poor growth and marked elongation responses seen with severe TE starvation (Fig. 2), the transient appearance of the regularly-shaped rod cell type seen in the optimally-growing cultures (+TE) was not clear during these -TE assays. This emphasizes the importance of using complete, TE-supplemented media for obtaining consistent results in the regular rod development assays, and offers a standardized way of performing TE-starvation when those cell types are required.

### Reversibility and specificity of the cell elongation response to TE-starvation

To investigate the reversibility of the TE-starvation phenotype in the Hv-Ca cultures, TEs were added at 15 h incubation after colony resuspension either by direct addition of the TE stock solution, or by dilution of the Hv-Ca culture (1:100) into Hv-Cab, Hv-Cab with 10x concentrated TE, or Hv-Ca (-TE) as a control. In all three media conditions containing additional TE, the cultures reverted to almost exclusively the plate cell morphotype at 12 h (Fig. 5). This is a similar duration required for the appearance of elongated cells upon colony resuspension in Hv-Ca (Fig. 4A, -TE), and shows that the effect of TE-starvation on morphology is readily reversible by resupply of the trace element nutrients. It is concluded that *H. volcanii* undergoes substantial cell elongation (and increased tubule development) in response to trace-element starvation, but not during starvation associated with stationary phase (+ TE) (Fig. 2–5, S2).

**Figure 5.**
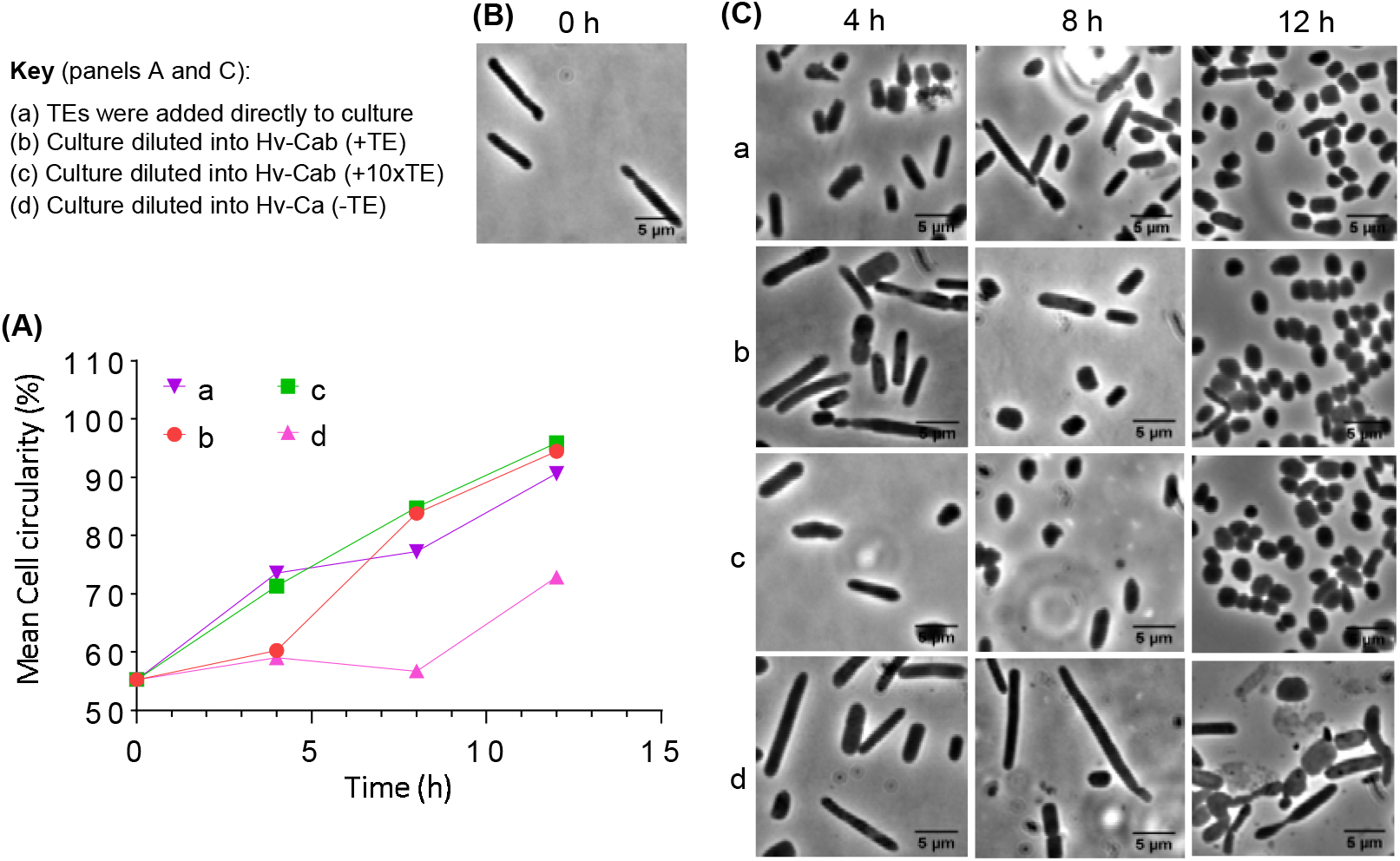
Reversal and specificity TE starvation. *H. volcanii* (H98 + pTA962) colonies from Hv-Ca agar were resuspended in Hv-Ca liquid medium. After 15 h of incubation, the culture was treated as indicated in the key, and then each new culture was monitored by sampling at 3 h intervals for microscopy over 12 hours. (A) The mean cell circularity (sample size N = 150 at each time point) versus time. (B) Representative phase-contrast image of the cells from the Hv-Ca (-TE) culture at 15 h, immediately prior to introducing TE. (C) Images containing cells after 4 h, 8 h and 12 h incubation under the conditions given in the key. Scale bars represent 5 microns.

The potential effects of individual components of the TE solution on *H. volcanii* cell shape were screened by testing a series of five TE solutions omitting either the Co, Cu, Fe, Mn or Zn. None of these individual ‘drop-out’ TE media matched the degree of cell elongation observed when all TE were omitted (Fig. S3), suggesting that the combined effect of multiple depleted elements causes the large change in cell shape seen in the -TE conditions. The Co and Fe drop-out media appeared to cause the most noticeable (although moderate) cell elongation response compared to the other metal drop-out media (Fig. S3).

### Microfluidic flow-chamber visualization of H. volcanii cellular development

We next sought to directly visualize morphological development during the early log growth phase in liquid culture. Cells resuspended from colonies were loaded into incubated microfluidic chambers for time-lapse microscopy. Time-series of rod-cell development in the media (+TE) are shown in Fig. 6A and 6B, and the complete time-series are available in Movies S1 and S4. These data show that plate cells can initially grow and divide as plates, and then most cells transition into rods as they grow and divide over several generations. We observed that rods persisted during incubation in the microfluidic culture system (particularly evident in Hv-YPCab), compared to the temporally quite narrow peak of rods seen in batch culture (Fig. 4A). The continual flow of fresh medium while cells are attached to the flow channels may therefore create signals that maintain rod morphology.

**Figure 6.**
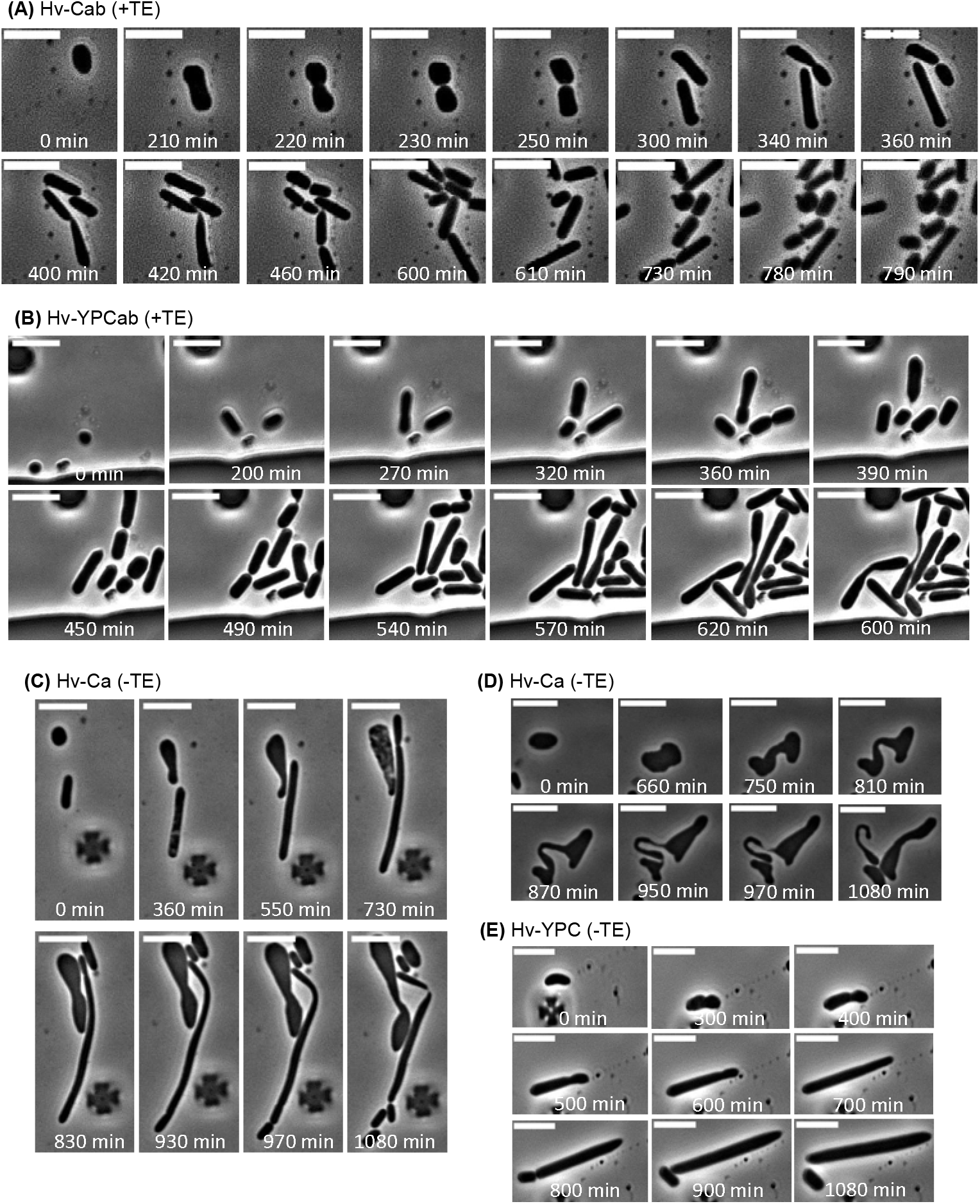
Time-lapse microscopy of *H. volcanii* (H98 + pTA962) with and without trace elements. Microfluidic chambers were loaded with cells and phase-contrast images were collected at 10 min intervals during perfusion (2 psi for 20 h). (A) Selected time frames showing initial growth after resuspension in Hv-Cab (+TE). (B) Selected time frames during microfluidic culture in Hv-YPCab (+TE). (C) Selected time frames showing the substantial cell elongation observed Hv-Ca (-TE) and a highly asymmetric division event. (D) Another field of cells cultured in Hv-Ca (-TE) showing slow growth and striking morphologies, including the generation of a cells that shows a distinctive highly curved narrowing or tubule. (E) Selected images during cell elongation in Hv-YPC (-TE).

Microfluidic cultures with media lacking the trace elements revealed striking morphological responses (Fig. 6C, 6D, 6E, and Movies S2, S3 and S5), including some extensively elongated cells exhibiting highly asymmetric division (Fig. 6C, 6E), and the generation of some bizarre irregular shapes (Fig 6D). This supported the batch culture results suggesting that formation of regularly shaped rods in early log phase is promoted in optimal growth conditions (in the presence of TE), whereas irregularly shaped and elongated shapes are associated with growth in the absence of TE and hence suboptimal growth conditions. Interestingly, tubules appeared to be generated by a combination of localized narrowing and elongation (tubulation), often in the mid-cell region of a cell, creating an approximate dumbbell-like appearance, followed by division near one end of the narrowing. We also saw that occasionally the narrow zone can re-widen, in a peristaltic-like wave along the cell (Movie S5).

### Requirement of tubulin-like protein CetZ1 for elongated shapes and tubule formation

In order to test whether the morphological changes require an internal cytoskeleton, we investigated the importance of the CetZ1 tubulin-like cytoskeletal protein in the cell shape changes. CetZ1 was previously found to be required for rod-cell development when cells become motile (6). A CetZ1-knock-out strain was subjected to the assays defined in Fig. 4, to determine if rod-shape or tubule development were affected under these conditions. In early-culture rod development assays, the Δ*cetZ1* strain failed to form rods at 10 h post-resuspension in Hv-Cab or Hv-YPCab, compared to clear rod formation in the wild-type control (Fig. 7A). Furthermore, both the elongated rods and the tubules observed in wild-type cells in the -TE conditions (Fig. 4) were not evident in Δ*cetZ1* cells (Fig. 7A). We also used Coulter cytometry to assess cell volumes of wild-type and CetZ1 knock-out strains. They showed very similar cell-size distributions in both media (15 h post-resuspension; Fig. 7C), despite the different shapes (Fig. 7A). However, both strains showed larger cells on average in Hv-Ca (-TE) than in Hv-Cab, demonstrating that cell size (regulation of cell division) also responds to trace-element starvation.

**Figure 7.**
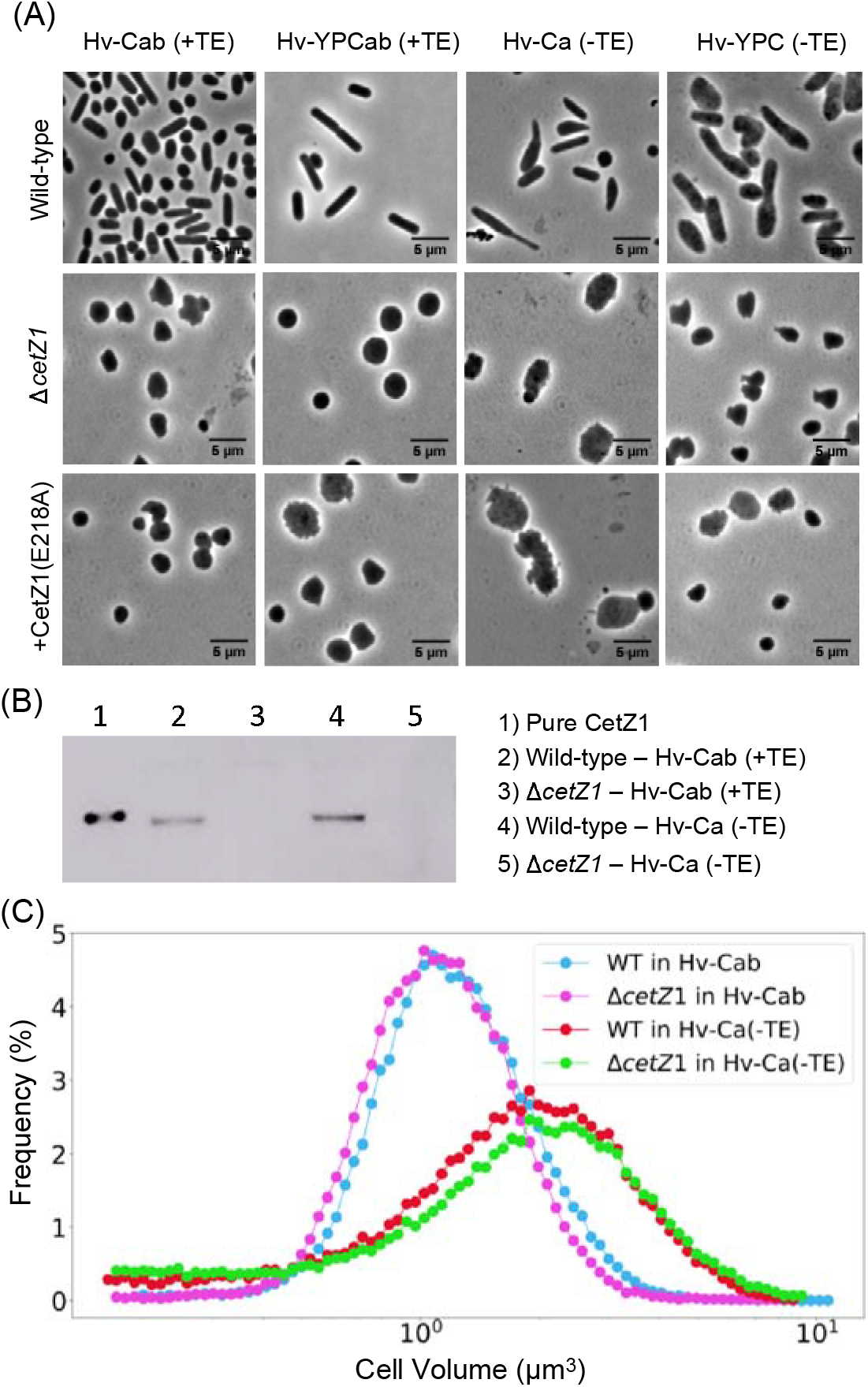
CetZ1 is essential for rod development during early-growth after colony resuspension and during TE-depletion. (A) Phase-contrast images of wild-type (H98 + pTA962), the CetZ1 knock-out (H98.Δ*cetZ1* (ID59) + pTA962), and the point-mutant (H98 + pTA962-*cetZ1*.E218A) were recorded 10 h post-resuspension of colonies in the indicated media (including 1 mM L-tryptophan, as the inducer for the *P.tna* promoter in pTA962). In Δ*cetZ1*, or during expression of CetZ1.E218A (-TE conditions), cells failed to form rods or substantially elongate, and sometimes exhibited a ruffled cell shape, showing small protrusions and grooves. (B) Western blot probing CetZ1 in the wild-type and CetZ1 knock-out strains at 15 h post-resuspension in the indicated media. (C) Coulter cytometry cell-volume distributions of the wild-type and CetZ1 knock-out strains at 15 h post-resuspension in Hv-Cab and Hv-Ca (-TE).29

We confirmed the expected absence of CetZ1 protein in the knock-out strain by Western blotting. We also tested whether the inclusion of TE influences cellular CetZ1 concentration, but observed no substantial difference (Fig. 7B). To verify that CetZ1 function is necessary for the rod and tubule formation, we expressed a dominant-inhibitory mutant (CetZ1.E218A) from a plasmid in a wild-type genetic background and then observed cell shapes. The E218A mutation in the GTPase active-site strongly inhibits CetZ1 function via hyper-stabilization of localized polymers, and was used previously for verification of CetZ1’s involvement in rod development in motility soft-agar (4). We found that expression of *cetZ1*.E218A blocked rod cell and tubule formation in the wild-type strain background (Fig. 7A), confirming its dominant influence on cell shape and the requirement for functional CetZ1 in trace-element-induced rod and tubule generation. Finally, we found that there were no detectable differences in growth rate or yield when comparing the wild-type, ΔcetZ1 and *cetZ1.E218A*-expression strains in Hv-YPC medium lacking TE (Fig. S4), showing that the morphology differences are not associated with substantial changes in growth during moderate TE limitation.

## Discussion

We report improved *H. volcanii* growth, in both rich and semi-defined media, by inclusion of eight supplementary trace elements (TE) as well as the identification and characterization of culture-dependent morphological changes that are more reproducible in the presence of TE.

Our surprising observation that TE-supplementation can improve growth and shape uniformity even in the complex medium containing yeast-extract made with high-purity salt, buffer, and water suggests that the optimal growth of *H. volcanii* requires the presence of relatively high levels of certain ‘trace’ metals. Previous *H. volcanii* media have been prepared with a trace elements solution containing Mn, Zn, Cu and Fe (13), but these are unlikely to represent all the minor elements utilized by *H. volcanii* (Fig. 3 and Fig. S3), and, furthermore, this four-element solution has only been widely adopted for use in minimal medium (10). More recent reports on trace-element supplementation in *H. volcanii* media have used six (14) of the elements used in the current study, and all eight elements have been used with synthetic medium (15), although in differing proportions. However, these studies did not report the influence these elements had on growth and morphology. Relatively high but variable metal ion concentrations occur in hypersaline lakes including the Dead Sea (the origin of *H. volcanii* DS2), which is especially rich in manganese (~100-150 μM) (1, 16).

Why would trace-element limitation trigger the development of *H. volcanii* elongation or tubules? Some of the factors that are related to morphological changes in bacteria are nutrient availabiliy, extracellular signalling, cell attachment and biofilm formation, motility/chemotaxis, predation and polar differentiation (17, 18). For example, cells, starving for multiple nutrients results in cell shape changes in a marine *Vibrio* species, where the pleomorphic cell populations stay viable for extended periods (19). In *Caulobacter crescentus*, phosphate starvation triggers stalk extension, which is thought to increase surface area for nutrient uptake (20, 21). Why *H. volcanii* changes cell shape during trace element starvation is not yet clear, however, the observation that these cells also get larger (Fig. 4B), thus reducing surface area-to-volume ratio, suggests that the overall response might do more than improving nutrient-uptake.

Our study also characterized the timing and conditions for *H. volcanii* rod development during the early stages of growth in dilute liquid cultures including supplemental TE. These rods were regularly-shaped and appear similar to the motile rods taken from soft agar (6). Indeed, early log cells were recently shown to be motile and suitable for direct observation of swimming (12). *H. volcanii* cell shape changes are also highly sensitive to factors other than nutrient availability; for example, we found that early-culture rod formation in the high-purity reagent medium, requires the presence of an *H. volcanii* plasmid, pTA962 (Fig. S1), which contains the endogenous pHV2 replication functions and marker genes (*pyrE2*, *hdrB*) that complement auxotrophic strains. A previous study noted a similar effect of the plasmid on the prevalence of rods (7), and we have also observed a slight decrease in cell size (unpublished data). While the plasmid does not appear to significantly affect growth rate (9), the results suggest that *H. volcanii* cell size and shape are influenced by auxotrophic status, plasmid replication and other functions, possibly through their impacts on metabolism. Other cellular responses, including the development of motility in soft agar, appear to be less sensitive to some of these effects (6). These findings emphasize the general importance of using strains that are genetically complemented to a wild-type status; simply supplying the required factors for auxotrophic growth is inadequate to avoid diverse effects on these cells.

The rod development that occurs during the early growth stages of a culture after resuspension or dilution of *H. volcanii* in fresh liquid medium is consistent with the sensing of nutrients or diluted biotic signals such as secreted quorum-sensing molecules. Consistent with this, we also observed cells retaining their rod shapes for extended periods when cells are exposed to a continuous flow of fresh medium. However, cells might also sense surface attachment and shear forces that differ between the flow chambers and liquid batch cultures. We also observed that cell shapes in agar surface colonies differed greatly from cell shapes in liquid cultures made with medium that was otherwise the same (both -TE), suggesting that the morphological effect of growth on agar (producing the plate morphotype) is dominant over the influence of trace element limitation, and could reflect sensing of different chemical or physical signals in the colonies. Since *H. volcanii* is capable of quorum sensing (22) and mechanosensing (23), further studies could ascertain how these processes might trigger changes in cell shape.

## Supporting information

Supplemental Results file

## Author contribution statement

Designed research: RTdS, MP, IGD. Contributed data: MFAH, MP (Fig. 1), RTdS (Fig. 2-7, S2-3, Movies S1-S5), DAP, HJB (Fig. S1, S4). Interpreted data: RTdS, MP, DAP, IGD. Wrote the manuscript: RTdS, IGD.

## Acknowledgements

This work was supported by the Australian Research Council (FT160100010 and DP160101076). M.P. and M.F.A.-H. were supported by the National Science Foundation grant 1817518. We are grateful for use of the equipment and support from the UTS Microbial Imaging Facility staff Louise Cole, Michael Johnson and Chris Evenhuis, and Solenne Ithurbide and Stephen Vadia for comments on the manuscript.

