## Supplemental Results file for "Improved growth and morphological plasticity of *Haloferax volcanii*"

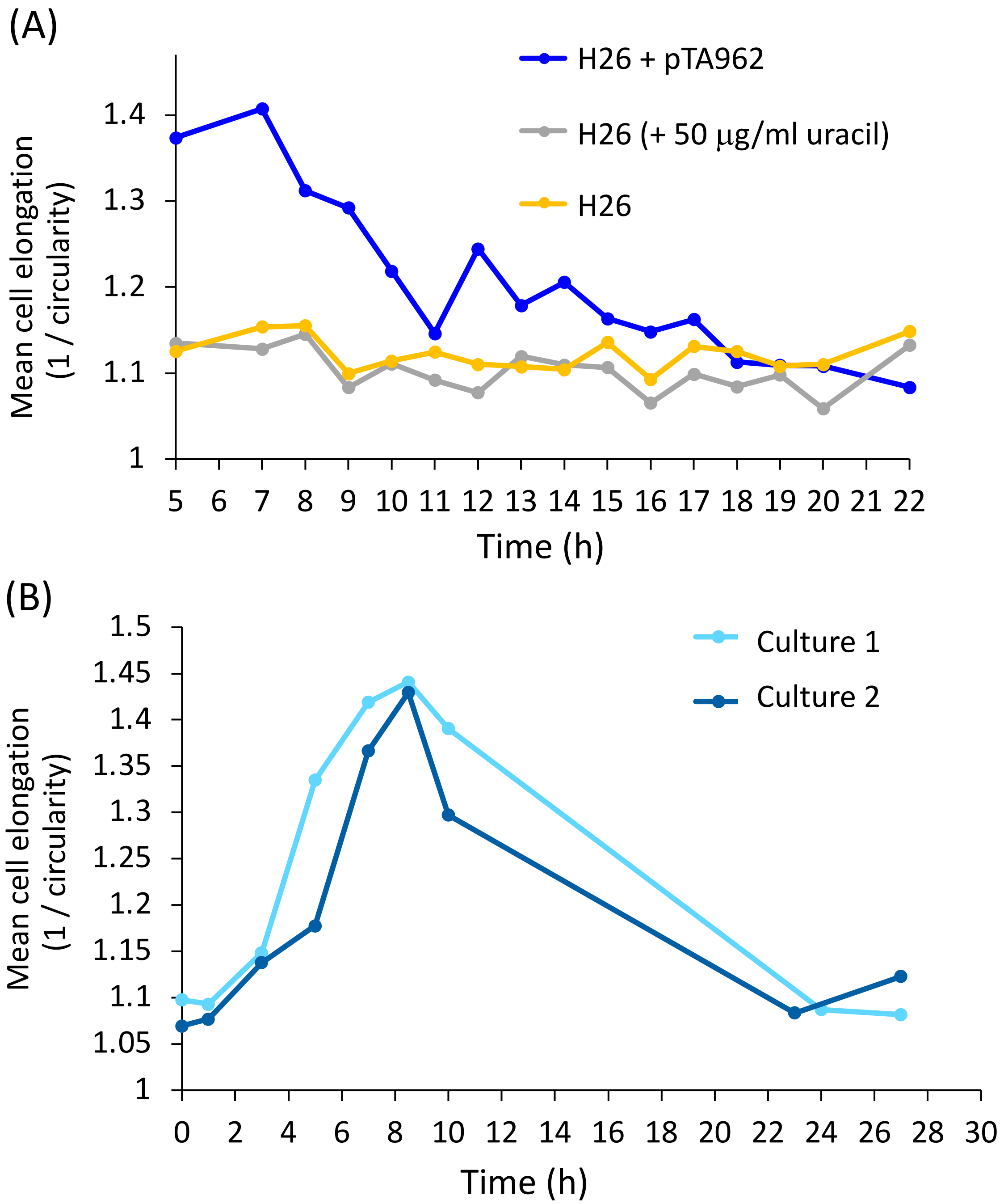


**Figure S1. (A) Effect of the pHV2-based plasmid (pTA962) on rod development in the early stage of culture.** The indicated *H. volcanii* strains were resuspended in liquid Hv-YPCab medium to $OD_{600\ nm}$ = 0.05 from fresh colonies grown on the respective agar medium. Cultures were incubated at 45°C at 200 rpm shaking, and samples were withdrawn at the indicated times and imaged with phase-contrast microscopy. Cell shapes were quantified as described in *Materials and Methods*. The peak of rod cells was observed only in the strain with pTA962. Supply of additional uracil (50 µg/ml uracil) into the Hv-YPCab medium (which already contains sufficient uracil for growth without supplementation) did not restore the rod phenotype in H26 (without plasmid). Each datapoint represents the mean cell elongation based on quantification of between 190 and 15,170 cells. **(B) Reproducibility of the timing of the peak of rod cell development, based on liquid starter cultures.** Two independent liquid starter cultures of *H. volcanii* H26 + pTA962 (grown at 45°C at 200 rpm shaking, $OD_{600\ nm}$ ~ 1.7) were used to inoculate fresh Hv-YPCab medium to $OD_{600\ nm}$ = 0.05, and then incubation was continued, and samples were taken, imaged and analysed as above. Each datapoint represents the mean cell elongation based on quantification of between 220-4187 cells.

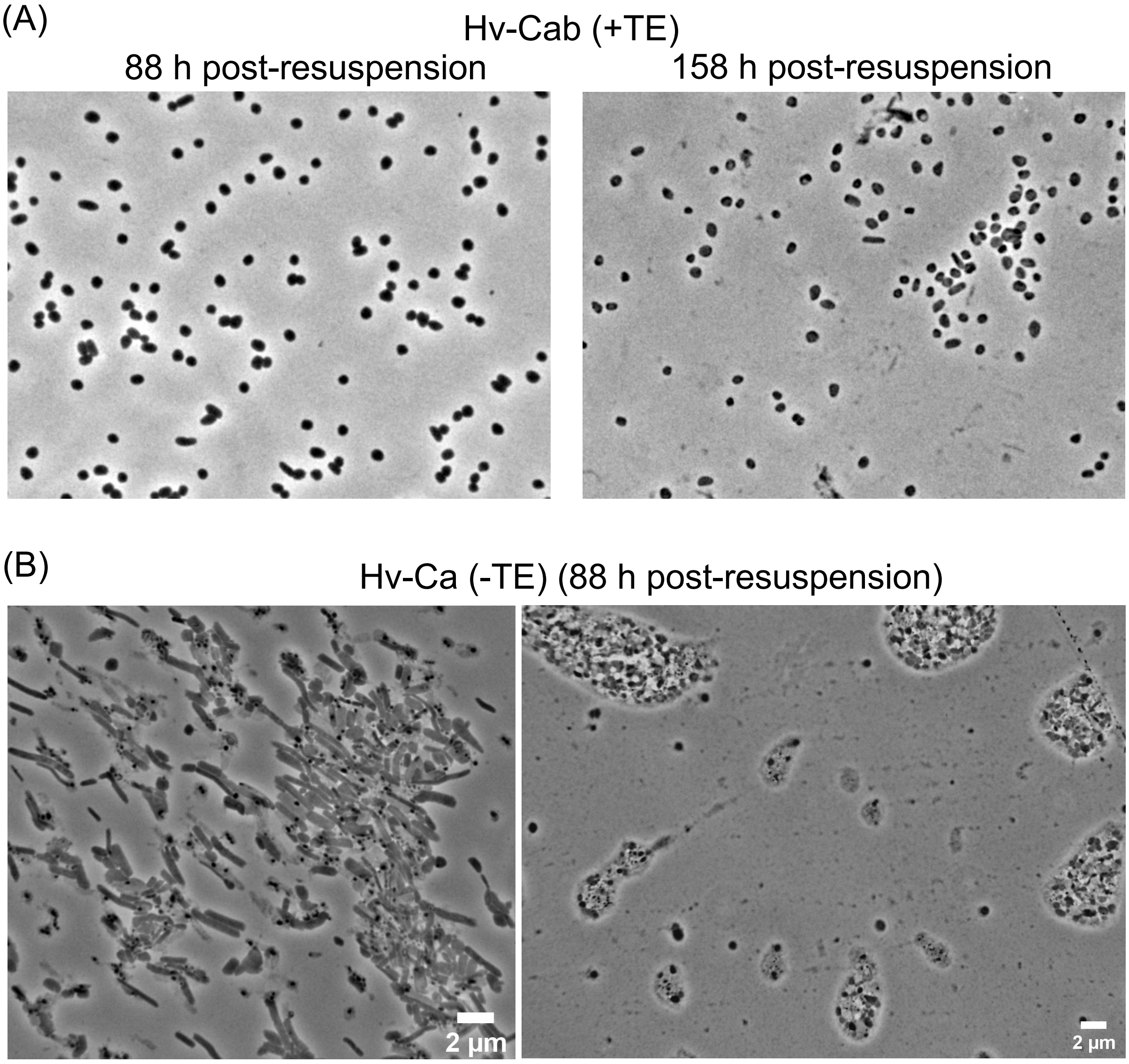


**Figure S2. Long-term stationary phase culture of *H. volcanii*.** Strain H98 (+ pTA962) colonies, grown on agar with (A) or without (B) supplementary trace elements in Hv-Ca medium, were resuspended (to $OD_{600}$ = 0.05) in the respective liquid media and cultured for the indicated times, before sampling for phase-contrast microscopy. (Refer to Fig. 4, for detailed analyses of the early stage of these conditions.) (A) With added trace elements, cell morphology remained almost exclusively the pleomorphic plate morphotype over extended time periods in stationary phase. (B) Without added trace elements, many late stationary phase cells show highly elongated morphology, phase-dark granules (left sample) and evidence of biofilm-like clustering (right sample) that became more obvious in longer-term cultures.

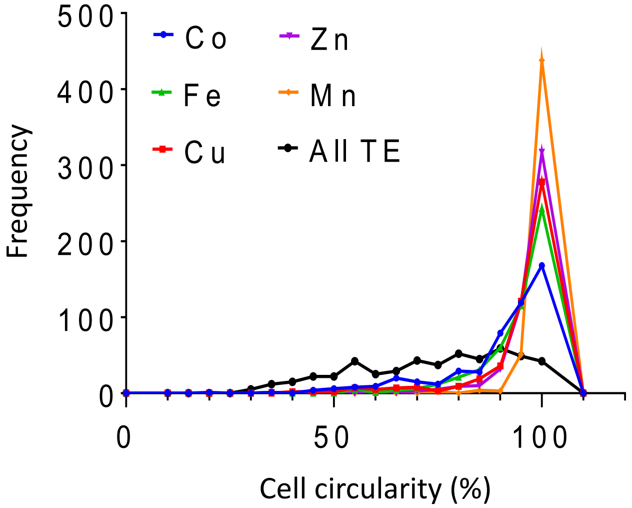


**Figure S3. Cell shape effects of individual omission of supplementary metals.** *H. volcanii* (H98 + pTA962) was resuspended in liquid Hv-Cab lacking supplemental Co, Fe, Cu, Zn, Mn or all TEs, from colonies grown on the same 'drop-out' agar medium. The cell circularity histograms are based on the indicated TE-dropout media, 15 h after inoculation into the liquid media (45°C at 200 rpm shaking), from two independent cultures (N = 500 cells). Omitting Co caused a slightly larger effect on cell elongation than omitting other metals, even though it is included at relatively low levels in the TE-solution (0.8 µM). *H. volcanii* cell shape could be especially sensitive to Co, or the residual concentration of Co could be extremely low in the media without added Co, relative to requirements, thereby eliciting a stronger response to its omission compared to other metals.

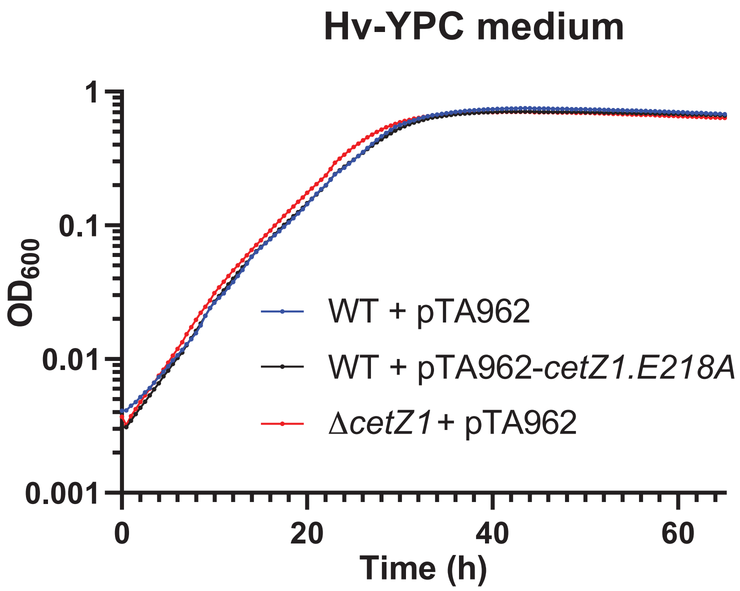


**Figure S4. Growth of *H. volcanii* and *cetZ1* mutant strains in Hv-YPC medium without supplementary TE.** Growth of *H. volcanii* (H98 + pTA962), as well as strains expressing the *cetZ1.E218A* dominant inhibitory mutant or carrying *cetZ1* deletion were grown into steady mid-log growth in Hv-YPC + 1 mM Trp, and then diluted into microtiter plates for monitoring growth ($OD_{600}$ over time) in the same medium, as described in the Methods. The data represent the mean of four independent cultures for each strain (each replicate culture was monitored simultaneously in three separate wells). These data show that the growth characteristics of the strains were indistinguishable.
